# Countries’ geographic latitude and their human populations’ cholesterol and blood pressure

**DOI:** 10.1101/308726

**Authors:** Yuhao Liu, Robert D. Brook, Xuefeng Liu, James Brian Byrd

## Abstract

**Background** Sunlight has been hypothesized to play a role in variation in cardiovascular disease according to geographic latitude. **Objectives** To evaluate the plausibility of sunlight as a factor in populations’ average cholesterol and blood pressure **Methods** We analyzed World Health Organization data including 180 or more countries’ age-standardized average cholesterol, age-standardized mean systolic blood pressure (BP), and age-standardized prevalence of raised BP, by geographic latitude, over decades. We also performed analysis by ultraviolet B light (UVB) exposure. **Results** Mean cholesterol increases with the distance of a country from the Equator. This relationship has changed very little since 1980. Similarly, in 1975, mean systolic BP and prevalence of raised BP were higher in countries farther from the Equator. However, the relationship between latitude and BP has changed dramatically; by 2015, the opposite pattern was observed in women. Countries’ average UVB exposure has a stable relationship with cholesterol over recent decades, but a changing relationship with BP. **Conclusions** Since sunlight exposure in a country is relatively fixed and its relationship with BP has changed dramatically in recent decades, countries’ average sunlight exposure is an unlikely explanation for contemporary country-level variation in BP. However, our findings are consistent with a putative effect of sunlight on countries’ average cholesterol, as well as a no longer detectable effect on BP decades ago. A parsimonious potential explanation for the relationship between light and cholesterol is that 7-dehydrocholesterol can be converted to cholesterol, or in the presence of ultraviolet light, can instead be converted to vitamin D.

## Introduction

In the 1980s, the incidence of cardiovascular disease was noted to be higher in countries farther from the Equator. (Fleck 1989) A popular hypothesis is that sunlight exerts a protective effect by reducing cholesterol and/or blood pressure. (Grimes et al. 1996; Patwardhan et al. 2017; Rostand 1997) A classic study in rabbits (Altschul 1953) and a more recent study in mice, (Geldenhuys et al. 2014) a cross-sectional study of humans, (Prodam et al. 2016) and a small clinical trial (Patwardhan et al. 2017) provide evidence of an effect of ultraviolet radiation on cholesterol. A parsimonious potential explanation is that 7-dehydrocholesterol can be converted to cholesterol, or in the presence of ultraviolet light, it can instead be converted to vitamin D. (Geldenhuys et al. 2014; Patwardhan et al. 2017) Sunlight’s effect on blood pressure is hypothesized to have a more complex biological basis, which might involve vitamin D, nitric oxide, and melanin. (Feelisch et al. 2010) There is experimental evidence to support an effect of ultraviolet B light on blood pressure. (Krause et al. 1998) Sunlight might also change cardiovascular risk factors by influencing countries’ agriculture, for example the availability of fresh fruits and vegetables. Whether effects of sunlight on cholesterol or blood pressure manifest at the level of global health is not well understood since most ecological studies relating sunlight exposure and/or latitude to cardiovascular risk factors are old and cross-sectional and involve relatively few countries.

New treatments for elevated cholesterol and blood pressure have been developed and disseminated since the 1970s, even as changes in health habits have swept the globe. It is possible that the relationship between latitude and cholesterol or blood pressure has changed accordingly, suggesting that sunlight is at most a small factor compared to other aspects of lifestyle. If instead, latitude or ultraviolet radiation has an unchanged relationship with a cardiovascular risk factor over decades despite lifestyle changes, a role for sunlight is more likely. Using decades of longitudinal data from over 180 countries, we examined the relationship between latitude and total cholesterol, and systolic blood pressure, with attention to changes over time. We also examined the relationship between ultraviolet B light exposure and countries’ cholesterol or blood pressure. In addition, we examined whether sex differences exist in these relationships.

## Methods

We obtained the country latitudes used in Google’s Public Data Explorer project. (Google) From the World Health Organization’s (WHO) Global Health Observatory, (World Health Organization) we obtained country-level age-standardized estimates of: mean total cholesterol (1980-2009), mean systolic blood pressure (1975-2015), and the prevalence of raised blood pressure (1975-2015). Raised blood pressure was defined by WHO as systolic blood pressure≥140 mm Hg or diastolic blood pressure≥90 mm Hg). Additional methods underlying the blood pressure measurements have been published. (NCD Risk Factor Collaboration 2017)

In addition, we obtained country-level ultraviolet B light exposure (averaged between 1997-2003) from the WHO Global Health Observatory. Some countries’ names were designated differently in the Google Public Data Explorer latitudes dataset and the WHO datasets. After harmonizing the country names, we used local regression (LOESS) plots to visualize the relationship between latitude and these endpoints. The R package ‘*ggplot2’* and its command ‘geom_smooth’ command were used to generate the LOESS curves and corresponding 95% confidence intervals. We also calculated Spearman correlation coefficients for distance from the Equator and countries’ mean cholesterol, mean systolic blood pressure, or the prevalence of raised blood pressure. We further evaluated whether our findings were stable over recent decades. Analyses were performed using R 3.4.3. (R Core Team (2017))

## Results

Latitude-based analyses included 188 countries’ mean cholesterol values, 180 countries’ mean systolic blood pressures, and 189 countries’ prevalences of raised blood pressure. Ultraviolet B radiation exposure-based analyses included 185 countries’ mean cholesterol values and 186 countries’ raised blood pressure prevalences.

Since at least the 1980s, average total cholesterol has been lowest in women (**Figure 1A**) and men (**Figure 1B**) living in countries at or near the Equator and has increased symmetrically with distance north or south of the Equator. Despite a downward shift in countries’ average cholesterol over recent decades, this symmetry around the Equator has been preserved. Absolute distance of a country from the Equator has correlated with total cholesterol in women (**Figure 1C**) and men (**Figure 1D**) since at least the 1980s (r_s_ = 0.66, *P*<2.2e^−16^ [females]; r_s_ = 0.67, *P*<2.2e^−16^ [males] in 1980; r_s_ = 0.51 *P*=1.2e^−13^ [females]; r_s_ = 0.57, *P*<2.2e^−16^ [males] in 2009).

**Figure 1.**
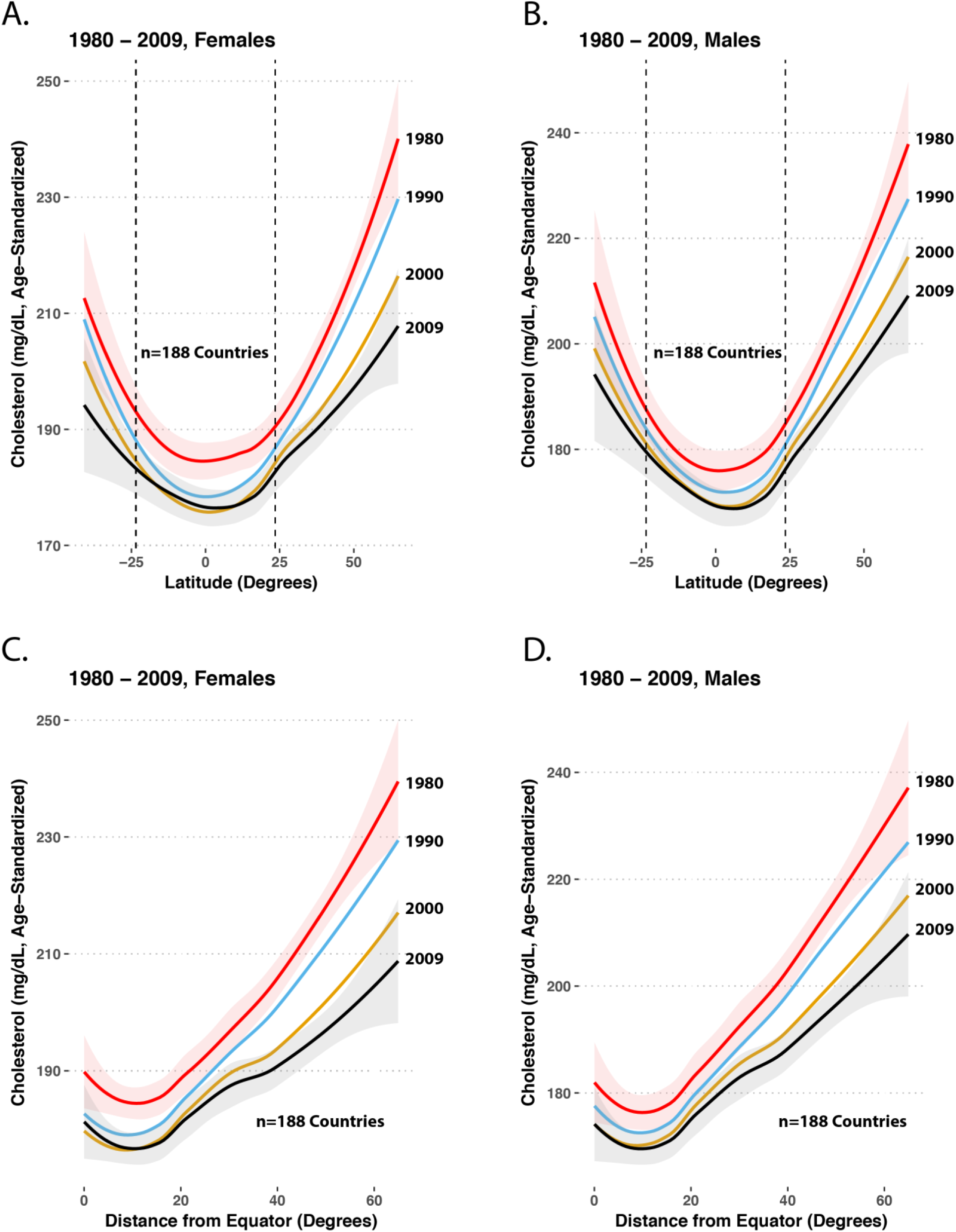
LOESS plots of latitude or distance from the Equator and countries’ mean total cholesterol (age-standardized). The top two panels show countries’ latitude and the mean cholesterol in females (**Panel A**) and males (**Panel B**). The vertical dotted lines represent 23.5 degrees North and South, the Tropics of Cancer and Capricorn, which define the tropics. The lower two panels show LOESS plots of absolute distance from the Equator in degrees and countries’ mean total cholesterol (age-standardized) in females (**Panel C**) and males (**Panel D**). To provide confidence intervals while maintaining visual clarity, the oldest and most recent years’ LOESS curves show the 95% confidence interval in pink and grey, respectively.

In 1975, countries’ mean systolic blood pressure followed a pattern similar to mean cholesterol, with systolic blood pressure increasing with distance north or south of the Equator in females (**Figure 2A**) and to a lesser extent in males (**Figure 2B**). As with cholesterol, countries’ mean systolic blood pressure exhibited symmetry north and south of the Equator. In contrast to cholesterol, mean systolic blood pressure’s relationship with latitude has been unstable over time. In fact, by 2015, the pattern had reversed itself in females (**Figure 2A**). In males, countries’ mean systolic blood pressure had a complex, bimodal relationship with latitude in 2015 (**Figure 2B**). Particularly in females, the changes in latitude’s relationship with age-standardized mean systolic blood pressure have been similar in the Northern and Southern hemispheres, preserving the north-south symmetry seen in the 1970s, with the Equator appearing as an inflection point. The slope of the relationship between absolute distance from the Equator and mean systolic blood pressure has been decreasing over time in females (**Figure 2C**)--in whom it has reversed direction in recent years to a negative relationship--and in males (**Figure 2D**, r_s_ = 0.52, *P*=4.0e^−14^ [females]; r_s_ = 0.49, *P*=2.9e^−12^ [males] in 1975; r_s_ = −0.28, *P*=0.0002 [females]; r_s_ = 0.07, *P*=0.32 [males] in 2015). Compared to mean systolic blood pressure, countries’ prevalences of raised blood pressure had similar LOESS plots (**Figures 3A and 3B**), but stronger symmetry around the Equator and marginally stronger correlation in the 1970s (r_s_ = 0.57, *P*<2.2e^−16^ [females]; r_s_ = 0.60, *P*<2.2e^−16^ [males] in 1975; r_s_ = −0.29, *P*=4.9e^−5^ [females], r_s_ = 0.13, *P*=0.09 [males] in 2015, **Figure 3**).

**Figure 2.**
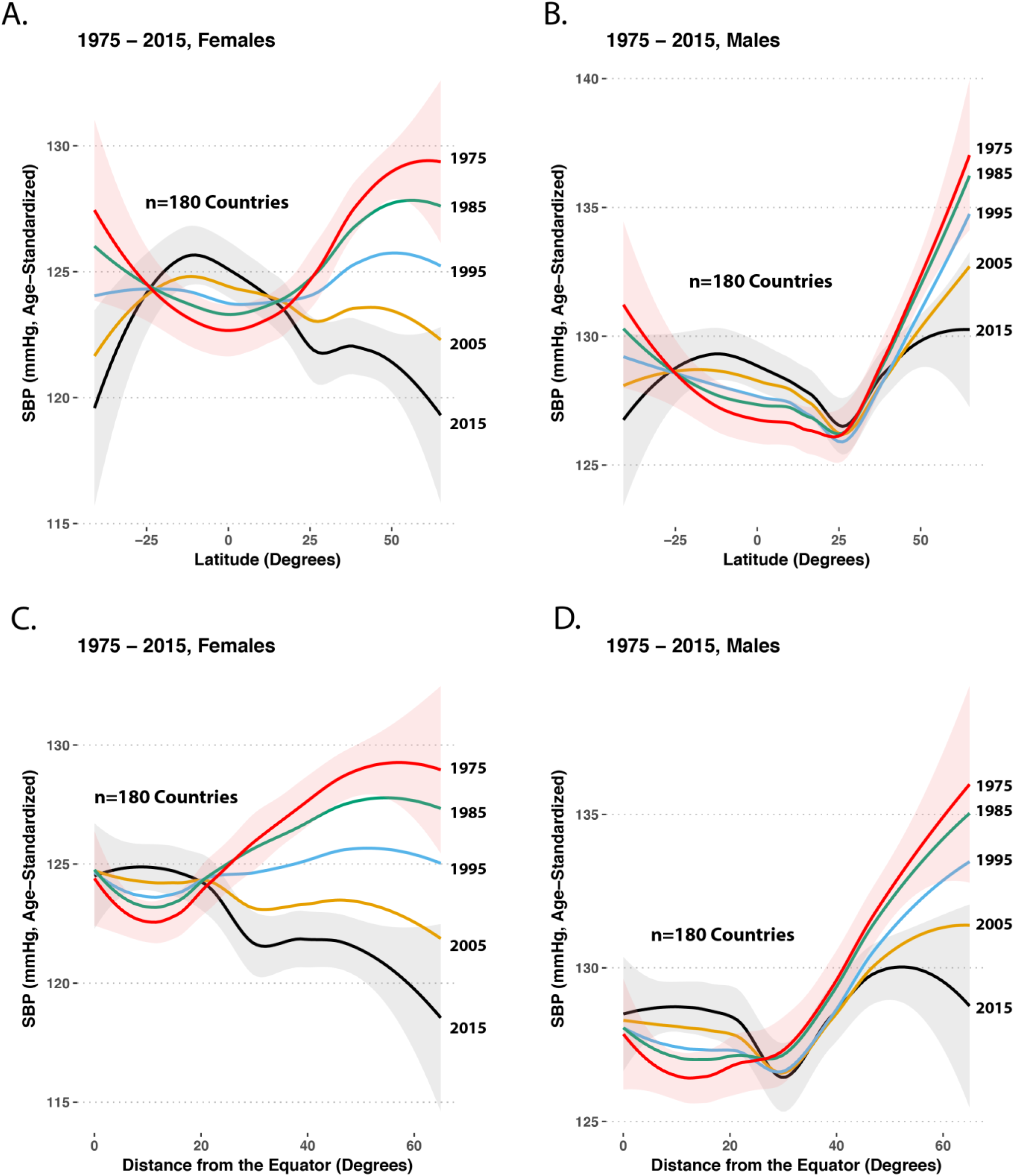
LOESS plots of latitude or distance from the Equator and countries’ mean systolic blood pressure (age-standardized estimate). The top two panels show mean systolic blood pressure according to latitude in females (**Panel A**) and males (**Panel B**), using data from every 10 years between 1975 and 2015. The lower two panels are LOESS plots of absolute distance from the Equator in degrees and countries’ mean systolic blood pressure (age-standardized estimate) in females (**Panel C**) and males (**Panel D**), respectively. To provide confidence intervals while maintaining visual clarity, the oldest and most recent years’ LOESS curves show the 95% confidence interval in pink and grey, respectively.

**Figure 3.**
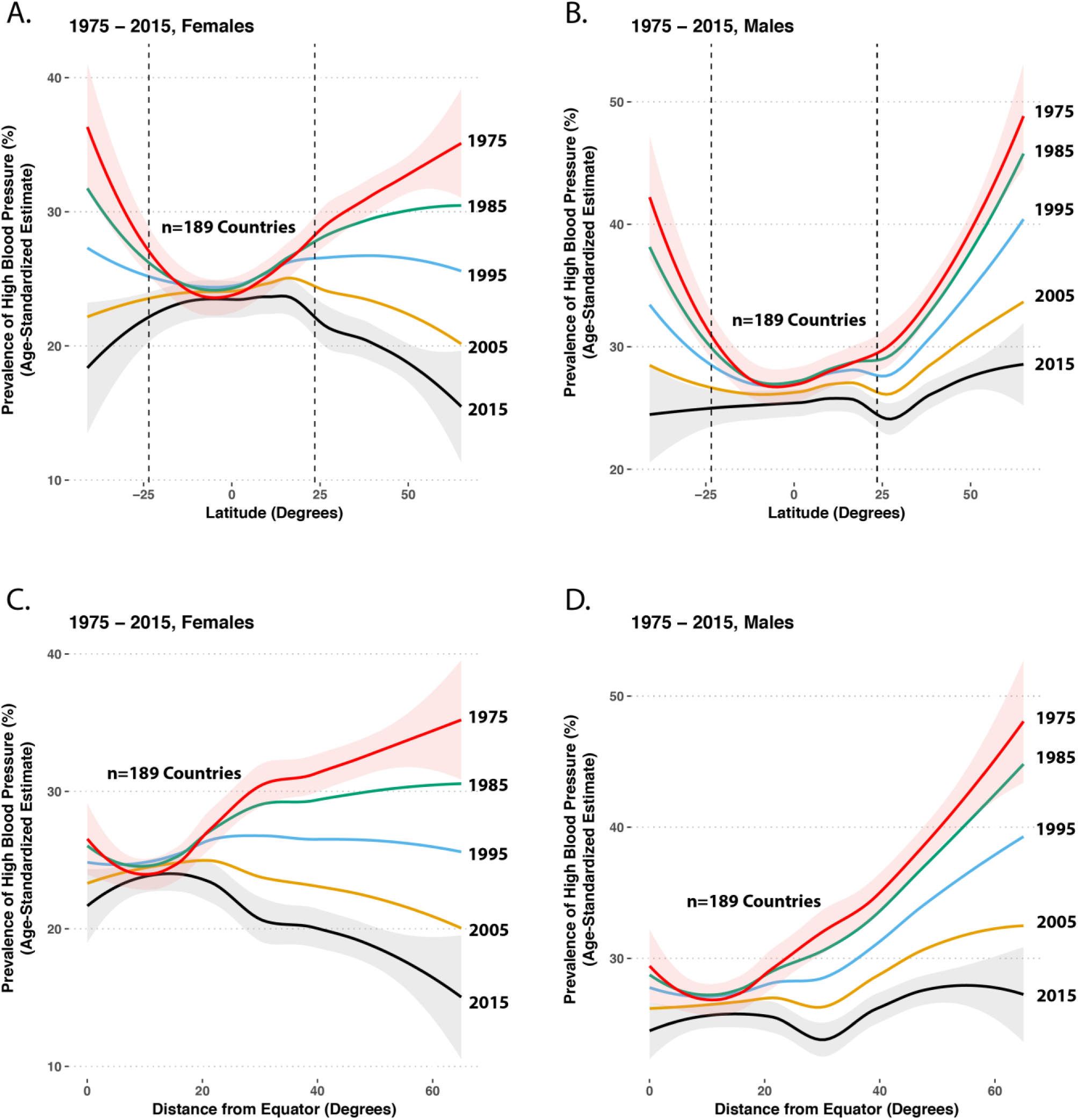
LOESS plots of latitude or distance from the Equator and countries’ prevalence of raised blood pressure (SBP≥140 or DBP≥90 [age-standardized estimate]). The upper two panels show the prevalence of raised blood pressure according to latitude in females (**Panel A**) and males (**Panel B**), using data from every 10 years between 1975 and 2015. The vertical dotted lines represent 23.5 degrees North and South, the Tropics of Cancer and Capricorn, which define the tropics. The lower two panels are LOESS plots of absolute distance from the Equator in degrees and countries’ prevalence of raised blood pressure in females (**Panel C)** and males (**Panel D**), respectively. To provide confidence intervals while maintaining visual clarity, the oldest and most recent years’ LOESS curves show the 95% confidence interval in pink and grey, respectively.

The relationship between ultraviolet B radiation and cholesterol or blood pressure mirrored their relationships with latitude. For decades, countries with higher average ultraviolet B radiation levels have had lower mean cholesterol in females (**Figure 4A**) and males (**Figure 4B**). As with latitude, the relationship between ultraviolet B radiation and raised blood pressure has been changing over time, with clear reversal of this relationship in females in 2015 compared to 1975 (**Figure 4C**) and dramatic changes in males (**Figure 4D**).

**Figure 4.**
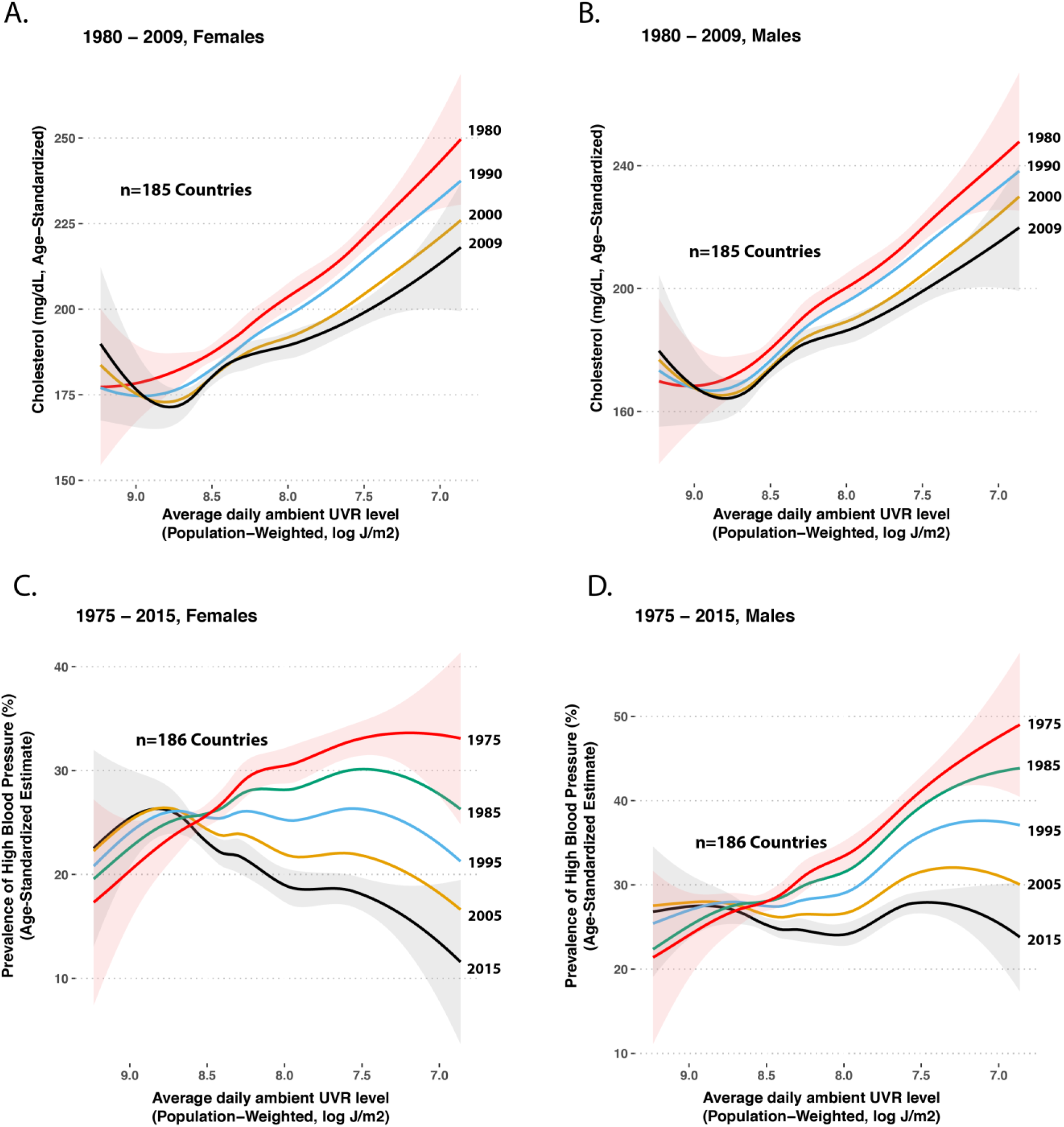
LOESS plots of countries’ log-transformed ultraviolet B radiation exposure (averaged between 1997-2003) and their populations’ mean cholesterol or prevalence of raised blood pressure (SBP≥140 or DBP≥90 [age-standardized estimate]). The upper two panels show mean cholesterol in females (**Panel A**) and males (**Panel B**) according to ultraviolet B radiation exposure. The lower two panels show LOESS plots of the relationship between log-transformed ultraviolet B radiation exposure and prevalence of raised blood pressure in females (**Panel C**) and males (**Panel D**). To provide confidence intervals while maintaining visual clarity, the oldest and most recent years’ LOESS curves show the 95% confidence interval in pink and grey, respectively.

## Discussion

The major new findings of this study are that for at least four decades, males’ and females’ average cholesterol levels have been higher in countries farther from the Equator, whereas mean systolic blood pressure and the prevalences of raised blood pressure showed a similar pattern decades ago, but no longer. In women, blood pressure’s relationship with latitude has reversed over time. Countries’ average ultraviolet B radiation has had a relatively constant relationship for decades with their populations’ mean cholesterol. The relationship between countries’ average ultraviolet B radiation exposure and blood pressure has changed dramatically over time.

In a small study (n=338) conducted at three sites in British Columbia, plasma cholesterol decreased with increasing latitude, a finding the authors attributed to differences in Rhesus Blood Group system. (Alfred et al. 1974) Conversely, hypertension prevalence or blood pressure have been reported to increase with distance north or south of the Equator or with increased solar radiation exposure within single countries (Cabrera et al. 2016; Rostand et al. 2016) and in studies of smaller groups of countries (Duranton et al. 2018; Rostand 1997) than we have analyzed. However, adjusting for vitamin D levels had no effect on the solar radiation-blood pressure relationship in a recent study, suggesting blood pressure’s variation by solar radiation levels is not mediated by vitamin D. (Rostand et al. 2016) Public availability of high-quality data facilitated our more comprehensive analysis of latitude’s and ultraviolet B radiation’s relationship with males’ and females’ cardiovascular risk factors estimated at the national level, around the globe and over decades. The focus of our study was to understand the relationship between latitude and cholesterol or blood pressure at the whole-globe level. Our findings contrast with the prior single-country analysis of latitude and cholesterol, suggesting a possible Simpson’s paradox, in which findings at a smaller scale of analysis are at odds with findings at a larger scale of analysis. Countries’ unique social history might explain such a paradox. Our latitude-blood pressure findings using data from decades ago are consistent with results from decades ago, but since then, the relationship not only has changed in men, it has reversed in women.

The factors explaining the stable relationship between countries’ latitude and their populations’ cholesterol are likely different from those explaining the relationship between latitude and blood pressure, which has changed dramatically over time. Exposure to sunlight has been proposed to explain the relationship between latitude and blood pressure, (Feelisch et al. 2010) but the marked changes in the relationship indicate it is no longer a key determinant of country-level differences in blood pressure, if it once was. Increased sunlight lowers cholesterol in experimental settings in rabbits and humans. (Altschul 1953; Patwardhan et al. 2017) Sunlight exposure could plausibly explain or contribute to the more stable relationship between latitude and cholesterol. An indirect effect of sunlight on cardiovascular risk factors through differences in agriculture is an alternate, plausible explanation of the relationship between UV light and cholesterol. There are likely other plausible explanations, as well. For example, countries’ gross domestic product varies by latitude. Interestingly, these differences in economic productivity might also be related to climate, (Masters and McMillan 2001) and thus related to sunlight.

Strengths of the current study include longitudinal analysis of high-quality data from more than 180 countries and the bringing together of multiple publicly available datasets to throw new light on an old question. The study is truly global in scale. The principal limitation of the study is directly related to this strength: ecological studies can show us what is true around the globe, but they permit only relatively weak inferences about how to explain what is seen. A second limitation is that our analysis compared countries’ UVB light exposure averaged between 1997-2003 to risk factors collected over a broader time span. Our analysis assumes stability of UVB light exposure, whereas there have been some regional changes in UVB light exposure due to the hole in the ozone layer. This issue affects only the UVB light analyses.

Ischemic heart disease (IHD) is the leading cause of death globally. A long-term 10% reduction in total cholesterol lowers risk of ischemic heart disease by 50% at age 40 and 20% at age 70, (Law et al. 1994) and a 20 mm Hg lower usual systolic blood pressure is associated with a 50% decrease in death from IHD and 50% decrease in death from stroke. (Lewington et al. 2002) As we seek new means of understanding cardiovascular risk reduction, additional studies to better understand the effect of sunlight on cardiovascular risk are needed.

## Conclusions

Since sunlight exposure in a country is relatively fixed and its relationship with BP has changed dramatically in recent decades, countries’ average sunlight exposure is an unlikely explanation for contemporary country-level variation in BP. However, our findings are consistent with a putative effect of sunlight on countries’ average cholesterol, as well as a no longer detectable effect on BP decades ago. A parsimonious potential explanation for the relationship between light and cholesterol is that 7-dehydrocholesterol can be converted to cholesterol, or in the presence of ultraviolet light, can instead be converted to vitamin D.

## Acknowledgements

JBB is funded by the National Institutes of Health award K23HL128909. We thank computational biologist Kasper Hansen for suggesting latitude might have a relationship with blood pressure after examining a map JBB had made. We thank Edward Roccella and Brahmajee Nallamothu for their thoughtful critical review of the manuscript.

## Notes

**Competing Financial Interests Delcaration:** The authors have no conflicts of interest.

## References

Alfred BM, Lee M, Desai ID. 1974. A relationship between plasma cholesterol level, latitude, and the rhesus blood group system. Hum Biol 46: 641–646.

Altschul R. 1953. Inhibition of experimental cholesterol arteriosclerosis by ultraviolet irradiation. N Engl J Med 249: 96–99.

Cabrera SE, Mindell JS, Toledo M, Alvo M, Ferro CJ. 2016. Associations of blood pressure with geographical latitude, solar radiation, and ambient temperature: Results from the chilean health survey, 2009–2010. Am J Epidemiol 183: 1071–1073.

Duranton F, Kramer A, Szwarc I, Bieber B, Gayrard N, Jover B, et al. 2018. Geographical variations in blood pressure level and seasonality in hemodialysis patients. Hypertension 71: 289–296.

Feelisch M, Kolb-Bachofen V, Liu D, Lundberg JO, Revelo LP, Suschek CV, et al. 2010. Is sunlight good for our heart? Eur Heart J 31: 1041–1045.

Fleck A. 1989. Latitude and ischaemic heart disease. Lancet 1: 613.

Geldenhuys S, Hart PH, Endersby R, Jacoby P, Feelisch M, Weller RB, et al. 2014. Ultraviolet radiation suppresses obesity and symptoms of metabolic syndrome independently of vitamin d in mice fed a high-fat diet. Diabetes 63: 3759–3769.

Google. Canonical latitudes for public data explorer project. Available: Available from https://developers.google.com/public-data/docs/canonical/countries_csv.

Grimes DS, Hindle E, Dyer T. 1996. Sunlight, cholesterol and coronary heart disease. QJM 89: 579–589.

Krause R, Buhring M, Hopfenmuller W, Holick MF, Sharma AM. 1998. Ultraviolet b and blood pressure. Lancet 352: 709–710.

Law MR, Wald NJ, Thompson SG. 1994. By how much and how quickly does reduction in serum cholesterol concentration lower risk of ischaemic heart disease? BMJ 308: 367–372.

Lewington S, Clarke R, Qizilbash N, Peto R, Collins R. 2002. Age-specific relevance of usual blood pressure to vascular mortality: A meta-analysis of individual data for one million adults in 61 prospective studies. Lancet 360: 1903–1913.

Masters WA, McMillan MS. 2001. Climate and scale in economic growth. Journal of Economic Growth 6: 167–186.

NCD Risk Factor Collaboration. 2017. Worldwide trends in blood pressure from 1975 to 2015: A pooled analysis of 1479 population-based measurement studies with 19.1 million participants. Lancet 389: 37–55.

Patwardhan VG, Mughal ZM, Padidela R, Chiplonkar SA, Khadilkar VV, Khadilkar AV. 2017. Randomized control trial assessing impact of increased sunlight exposure versus vitamin d supplementation on lipid profile in indian vitamin d deficient men. Indian J Endocrinol Metab 21: 393–398.

Prodam F, Zanetta S, Ricotti R, Marolda A, Giglione E, Monzani A, et al. 2016. Influence of ultraviolet radiation on the association between 25-hydroxy vitamin d levels and cardiovascular risk factors in obesity. J Pediatr 171:83–89.e81.

R Core Team (2017). R: A language and environment for statistical computing. R Foundation for Statistical Computing, Vienna, Austria.

Rostand SG. 1997. Ultraviolet light may contribute to geographic and racial blood pressure differences. Hypertension 30: 150–156.

Rostand SG, McClure LA, Kent ST, Judd SE, Gutierrez OM. 2016. Associations of blood pressure, sunlight, and vitamin d in community-dwelling adults. J Hypertens 34: 1704–1710.

World Health Organization. Global health observatory data repository. Available: Available from http://apps.who.int/gho/data/node.imr.

